# Recurrent *PD-L1* Structural Rearrangements in Natural Killer/T Cell Lymphoma Patients with Complete Response to PD-1 Blockade Therapy

**DOI:** 10.1101/372383

**Authors:** Jing-Quan Lim, Tiffany Tang, Qing-qing Cai, Daryl Tan, Maarja-Liisa Nairismägi, Yurike Laurensia, Burton Kuan Hui Chia, Rou-Jun Peng, Jabed Iqbal, Da Chuan Huang, Tammy Song, Wan Lu Pang, Daryl Ming Zhe Cheah, Cedric Chuan Young Ng, Vikneswari Rajasegaran, Huangming Hong, Eric Tse, Benjamin Mow, Qi Chun Cai, Li-Mei Poon, Jing Tan, Nicholas Francis Grigoropoulos, Yeow Tee Goh, Colin Phipps, Olaf Rötzschke, Chee Leong Cheng, Yuh Shan Lee, Yvonne Loh, Miriam Tao, Mohamad Farid, Rex Au-Yeung, Thomas Sau-Yan Chan, Siok-Bian Ng, Yok-Lam Kwong, William Hwang, Wee-Joo Chng, Thomas Tousseyn, Patrick Tan, Bin Tean Teh, Chiea Chuen Khor, Steve Rozen, ICGC Blood Cancer T-cell and NK-cell lymphoma group, Jin-Xin Bei, Tongyu Lin, Soon Thye Lim, Choon Kiat Ong

**Author notes:** Drs Jing Quan Lim, Tiffany Tang, Qing-qing Cai and Daryl Tan contributed equally to this work. Corresponding Authors: Dr. Choon Kiat ONG, Principal Investigator, Lymphoma Genomic Translational Research Laboratory, Division of Medical Oncology, National Cancer Centre Singapore, 11 Hospital Drive, 169610, Singapore; Prof. Soon Thye LIM, Head, Division of Medical Oncology, National Cancer Centre Singapore, 11 Hospital Drive, 169610, Singapore; Prof. Tongyu LIN, Department of Medical Oncology, Sun Yat-sen University Cancer Center, Guangzhou 510060, China; Prof. Jin-Xin BEI, Principal Investigator, State Key Laboratory of Oncology in South China, Sun Yat-sen University Cancer Center, Guangzhou 510060, China.

## Abstract

This study aims to identify recurrent genetic alterations in relapsed or refractory (RR) natural-killer/T-cell lymphoma (NKTL) patients who have achieved complete response (CR) with programmed cell death 1 (PD-1) blockade therapy. Seven of the eleven patients treated with pembrolizumab achieved CR while the remaining four had progressive disease (PD). Using whole genome sequencing (WGS), we found recurrent clonal structural rearrangements (SR) of the *PD-L1* gene in four of the seven (57%) CR patients’ pretreated tumors. These *PD-L1* SRs consist of inter-chromosomal translocations, tandem duplication and micro-inversion that disrupted the suppressive function of *PD-L1* 3’UTR. Interestingly, recurrent *JAK3*-activating (p.A573V) mutations were also validated in two CR patients’ tumors that did not harbor the *PD-L1* SR. Importantly, these mutations were absent in the four PD cases. With immunohistochemistry (IHC), PD-L1 positivity could not discriminate patients who archived CR (range: 6%-100%) from patients who had PD (range: 35%-90%). PD-1 blockade with pembrolizumab is a potent strategy for RR NKTL patients and genomic screening could potentially accompany PD-L1 IHC positivity to better select patients for anti-PD-1 therapy.

## Main Text

## Introduction

Natural-killer/T cell lymphoma (NKTL) is an uncommon and aggressive malignancy with a predilection for Asian, Mexican and South American populations (*1*). With the exception of Japan, it is the most common mature T-cell lymphoma in Asia (*2*). Neoplastic cells are invariably infected by the Epstein Barr virus (EBV) and are characterized by a cytotoxic phenotype (*3*).

In recent years, immune checkpoint (ICP) inhibitors have shown promising objective response rates (ORR) in the treatment of many malignancies (*4*). Of note, the most impressive result is 80% ORR from the use of programmed death-1 (PD-1 or CD279) inhibitors in relapsed or refractory (RR) Hodgkin lymphoma (HL) (*5, 6*). Currently, clinical studies involving non-small-cell lung carcinoma, melanoma and bladder cancer have generally concluded that immunohistochemistry (IHC) positivity of programmed death-ligand 1 (PD-L1) coincides with greater likelihood of response to PD-1/PD-L1 blockade. Intriguingly, there was also a lower but definite response rate in patients with PD-L1-negative tumors (*7*). These observations suggest that more information could be harnessed from the tumors and augment the current de facto criteria of selecting patients for PD-1 blockade therapy (*8, 9*).

With the major advancements in sequencing technologies, recurring somatic mutations altering the JAK-STAT pathway, epigenetic modifiers, *DDX3X* gene and germline genetic predisposition in the *HLA-DPB1* gene have been found in NKTL patients but none of these studies have investigated these tumors using whole genome sequencing (WGS) techniques (*10-16*). In order to explore the NKTL genome for actionable alterations associated to PD-1 blockade therapy, we employed WGS data to study the somatic alterations of 11 pretreated NKTL tumors that have corresponding clinical response data to pembrolizumab treatment.

## Results

### Exceptional response to pembrolizumab in relapsed or refractory NKTL patients

Eleven NKTL patients from Singapore, China and Hong Kong who were relapsed or refractory (RR) to L-asparaginase containing chemotherapy regimens, after a median of two (range: 1-5) lines of treatments, were included into this study (Table 1). These 11 pembrolizumab-treated patients had a median age of 42 years old at diagnosis (range: 27-66 years) and a median follow-up of 11 months (range: 2-25 months) since treated with pembrolizumab. Sixty-four percent (7 of 11 cases) of the patients achieved complete responses (CR) while 36% (4 of 11 cases) of the patients had progressive disease (PD). Two patients (NKTL26 & NKTL31) remained in remission from pembrolizumab for more than two years, which is rare in RR NKTL (*17*). The most recent pembrolizumab-treated case (NKTL28) also achieved ongoing remission for at least 6 months. The median duration of response to pembrolizumab (for responding patients) was 14 months.

**Table 1.** Clinical information and outcomes before and after pembrolizumab treatment of 11 NK/T-cell lymphoma patients.

|  | Case | Sex | Age, yr | Stage | Elevated LDH | IPI<sup>1</sup> | ECOG | OS, mth | PFS, mth | Treatments prior to Pembrolizumab |  |  |  | Pembrolizumab treatment |  |
| --- | --- | --- | --- | --- | --- | --- | --- | --- | --- | --- | --- | --- | --- | --- | --- |
|  |  |  |  |  |  |  |  |  |  | CTx (cycles) | RT | TP | PD-L1+, % (H-score) | Outcome | DOR<sup>2</sup>, mth |
| Complete Responders (CR) | NKTL1 | M | 49 | IV | N | 2 | 1 | 49+ | 19 | GELOX (4), SMILE (5), Romidepsin+Bortezomib, BV+Benda, Lenalidomide+Dara | Nil | Nil | 100% (250) | CR: PET/CT<br>EBV DNA: negative then became positive and remained stable | 20+ |
|  | NKTL26 | M | 32 | I | Y | 1 | 1 | 44+ | 2 | SMILE (2), Vinc+DXM+Lasp (1), GELOX (6) | Yes | Nil | 40% (35) | CR: PET/CT<br>EBV DNA: ND | 24+ |
|  | NKTL28 | M | 46 | IV | Y | 4 | 3 | 9+ | 0 | SMILE (2), P-GEMOX (1) | Nil | Nil | 70% (190) | CR: PET/CT<br>EBV DNA: negative | 6+ |
|  | NKTL29 | M | 48 | I | N | 0 | 0 | 13+ | 4 | Ifos+MTX+VP-16+DXM+Pasp (4) | Nil | Nil | 6% (7) | CR: PET/CT<br>EBV DNA: negative | 9+ |
|  | NKTL30 | M | 38 | IV | Y | 4 | 3 | 19+ | 6 | SMILE (5) | Nil | Nil | 60% (120) | CR: PET/CT<br>EBV DNA: negative | 11+ |
|  | NKTL31 | M | 27 | IV | Y | 4 | 0 | 67+ | 17 | Lasp+DXM+Vinc+AraC (4), CHOP (2), P-GEMOX (2), DXM+Pasp+mitoxantrone+VP-16 (4) P-GEMOX+VP-16 (2) | Nil | Auto-HSCT with BEAM + Thalidomide (maintenance) | 20% (20) | CR: CT & MRI<br>EBV DNA: negative | 24+ |
|  | NKTL43 | M | 29 | IV |  |  | 2 | 116 | 73 | m-BACOD (4), PIGLET (5), SMILE (3) | Yes | Nil | 90% (190) | CR: PET/CT<br>EBV DNA: negative<br>Patient subsequently underwent MUD BMT and died from GVHD. | 14 |
| Non-CR | NKTL25 | M | 30 | IV | N | 2 | 0 | 14 | 10 | SMILE (6), GEMOX (1) | Yes | Allo-HSCT | 72% (126) | PD: DOD | NA |
|  | NKTL27 | M | 59 | IV | Y | 2 | 0 | 19 | 2 | SMILE (3), GIFOX (4) | Nil | Nil | 50% (85) | PD: DOD | NA |
|  | NKTL44 | M | 66 | IV |  |  | 1 | 37 | 21 | SIMPLE (6) | Nil | Nil | 90% (170) | PD: DOD | NA |
|  | NKTL45 | M | 42 | IV |  |  | 1 | 94 | 87 | SMILE (6), GEMOX (1) | Nil | Allo-HSCT | 65% (70) | PD: DOD | NA |

<sup>1</sup>IPI: International Prognostic Index
<sup>2</sup>DOR: Durability of response as of Jan 2018; + indicates ongoing survival
BV, brentuximab vedotin; Benda, bendamustine; Dara, daratumumab; Vinc, vincristine; DXM, dexamethasone; Lasp, L-asparaginase; Ifos, ifosfamide; MTX, methotrexate; VP-16, etoposide; Pasp, Peg-Lasparaginase; AraC, cytarabine; ND, not done; RT, radiotherapy; TP, transplant

### PD-L1 positivity could not stratify response to pembrolizumab in NKTL patients

To verify if PD-L1 positivity in NKTL tumors could predict response to pembrolizumab, we proceeded to determine the positivity of PD-L1 in all 11 pretreated NKTL tumors using IHC. The same pathologist assessed PD-L1 positivity in all the tumors in this study to ensure consistency (Table 2). The PD-L1 positivity in the tumor cells varied greatly in both patients who achieved CR and PD. PD-L1 positivity in the pretreated tumors of the patients with CR ranged from 6% to 100% while the PD-L1 staining intensity among patients with PD ranged from 35% to 90%. Hence PD-L1 staining intensity could not differentiate between patients who achieved CR and those who had PD. Interestingly, NKTL29 had only 6% of tumor cells stained positive for PD-L1 but achieved CR from pembrolizumab. Apart from this PD-L1^low^ CR case, all four PD cases were strongly stained for PD-L1, with an average of 69% (range: 50% - 90%) tumor cells stained positively for PD-L1. This is concordant with clinical trials reporting that antitumor activity from PD-1 blockade therapy was also observed in melanoma and non small-cell lung carcinoma patients with low baseline PD-L1 positivity (*18, 19*). Our current study shows that some patients with low PD-L1 positivity may have good responses to PD-1 blockade.

**Table 2.** Membraneous PD-L1 immunohistochemical staining grade, PD-L1 H-score and PD-L1 positivity cells in the pretreated NKTL tumors of our study cohort.

| Sample ID | Response to pembrolizumab | *PD-L1* Rearranged | PD-L1 positivity %, strongest stain grade | PD-L1 lymphocytes 1+ <sup>a</sup> | PD-L1 lymphocytes 2+ <sup>a</sup> | PD-L1 lymphocytes 3+ <sup>a</sup> | H-score for PD-L1 |
| --- | --- | --- | --- | --- | --- | --- | --- |
| NKTL1 | CR | Yes | 100%, 3+ | 0% | 50% | 50% | 250 |
| NKTL26 | CR | Yes | 35%, 2+ | 30% | 5% | 0% | 40 |
| NKTL28 | CR | Yes | 70%, 3+ | 0% | 20% | 50% | 190 |
| NKTL31 | CR | Yes | 20%, 1+ | 20% | 0% | 0% | 20 |
| NKTL29 | CR | No | 6%, 2+ | 5% | 1% | 0% | 7 |
| NKTL30 | CR | No | 60%, 3+ | 10% | 40% | 10% | 120 |
| NKTL43 | CR | No | 90%, 3+ | 20% | 40% | 30% | 190 |
| NKTL25 | PD | No | 72%, 3+ | 20% | 50% | 2% | 126 |
| NKTL27 | PD | No | 50%, 3+ | 20% | 25% | 5% | 85 |
| NKTL44 | PD | No | 90%, 3+ | 20% | 60% | 10% | 170 |
| NKTL45 | PD | No | 65%, 2+ | 60% | 5% | 0% | 70 |

<sup>a</sup> Stain grade: 0, no; 1+, weak; 2+, moderate; 3+, strong. CR, complete response; PD, progressive disease.

### Whole genome sequencing and analysis of 11 RR NKTL pembrolizumab-treated patients

To identify genomic biomarkers of response to PD-1 blockade therapy in NKTL, whole genome sequencing was performed on pairs of tumor-normal samples obtained from 11 patients who were subsequently treated with pembrolizumab. The NKTL tumors and, whole blood or buccal swabs, were sequenced to an average depth of 66.6x and 37.5x, respectively (Supplementary Table 1). Somatic variant calling yielded an average of 1.15 single nucleotide variants (SNVs) and microIndels per Mb for each paired sample. An average of 39 (range: 1 – 80, Supplementary Table 2) somatic non-silent protein-coding variants per sample was identified and is comparable to previous reports on whole-exome sequencing of fresh-frozen NKTL samples (range: 41-42) (*10, 11*). In total, we found 10 genes to be recurrently mutated (Supplementary Fig. 1). Among them, only *PD-L1* structural rearrangement (SR) and *JAK3*-activating mutations (p.A573V) were recurrent and mutually exclusive to one another among the initial tumors of patients who achieved CR. Furthermore, *PD-L1* SR is the most frequent somatic alteration identified in four of seven (57%) initial tumors of patients who achieved CR (Fig. 1A). These *PD-L1* SRs consist of inter-chromosomal translocations (NKTL28 & NKTL31), tandem duplication (NKTL26) and micro-inversion (NKTL1) that disrupted the 3’UTR of *PD-L1* (Fig. 1B). In NKTL28 and NKTL31, exon 7 of *PD-L1* was translocated to 2q24.2 and intron 6 of *PD-L1* was translocated to 6p12.2, respectively (Supplementary Fig. 2). In NKTL26, the right breakpoint of tandem duplication was located within the 3’UTR of *PD-L1* and the left breakpoint was validated to be ∼32kbp upstream. This duplication event yielded a copy of 3’UTR-disrupted and wild type copy of *PD-L1* (Supplementary Fig. 3). The final *PD-L1* SR in NKTL1 consists of a 206 bp micro-inversion that sits entirely within the 3’UTR of *PD-L1*. These somatic alterations were absent in the initial tumors from the four patients who had PD with pembrolizumab.

**Fig. 1:**
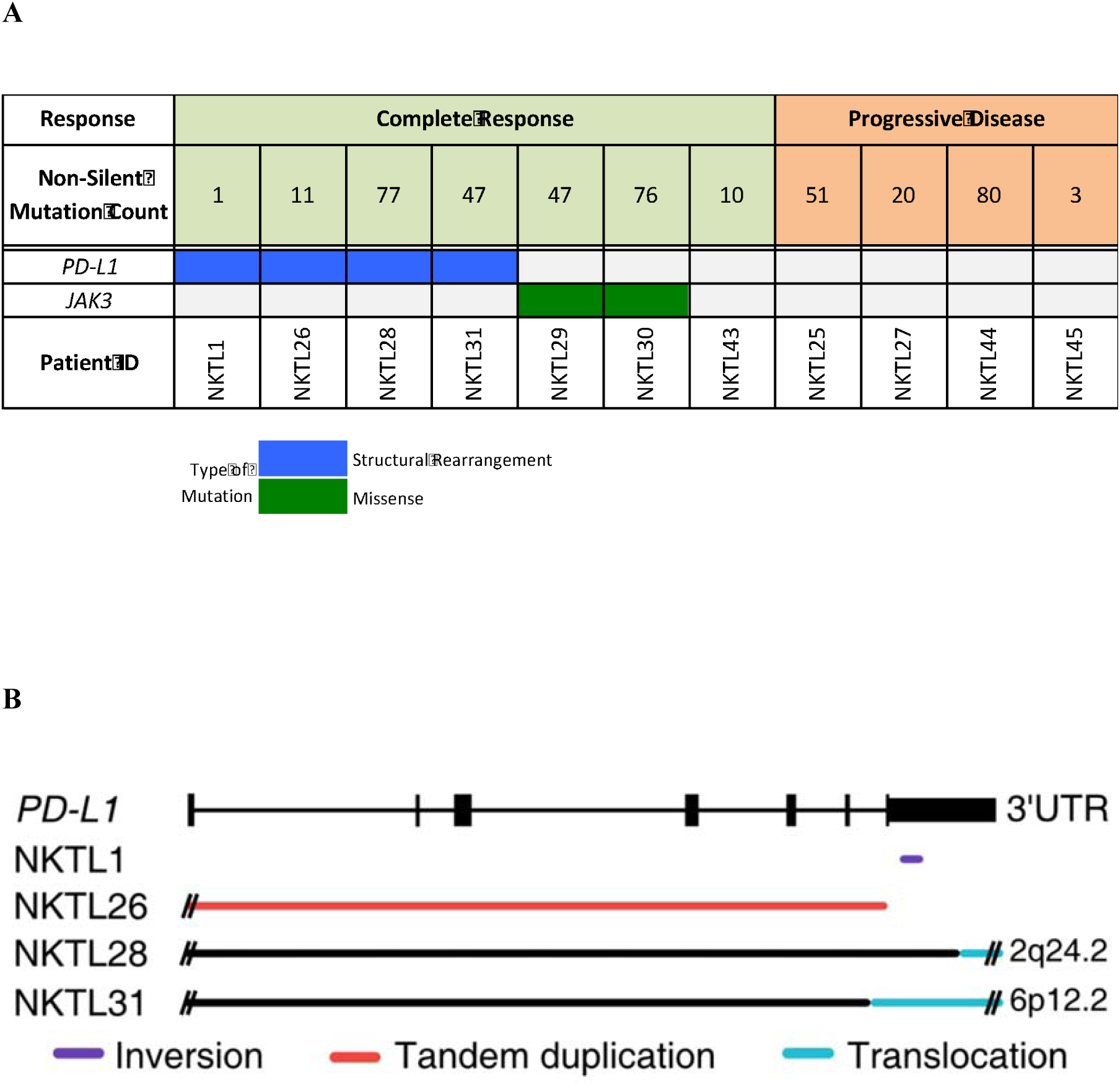
Genomic profiles of 11 pretreated NKTL tumors from patients who were subsequently treated with pembrolizumab. (A) Staircase plot of recurrent and mutually exclusive non-silent genomic alterations found in the 11 pairs of NKTL-normal whole-genome sequencing data. The top of the staircase plot denotes the number of non-silent mutations. SG: Singapore, HK: Hong Kong. (B) Schematics of the *PD-L1* structural rearrangements that were validated in our study.

**Fig. 2:**
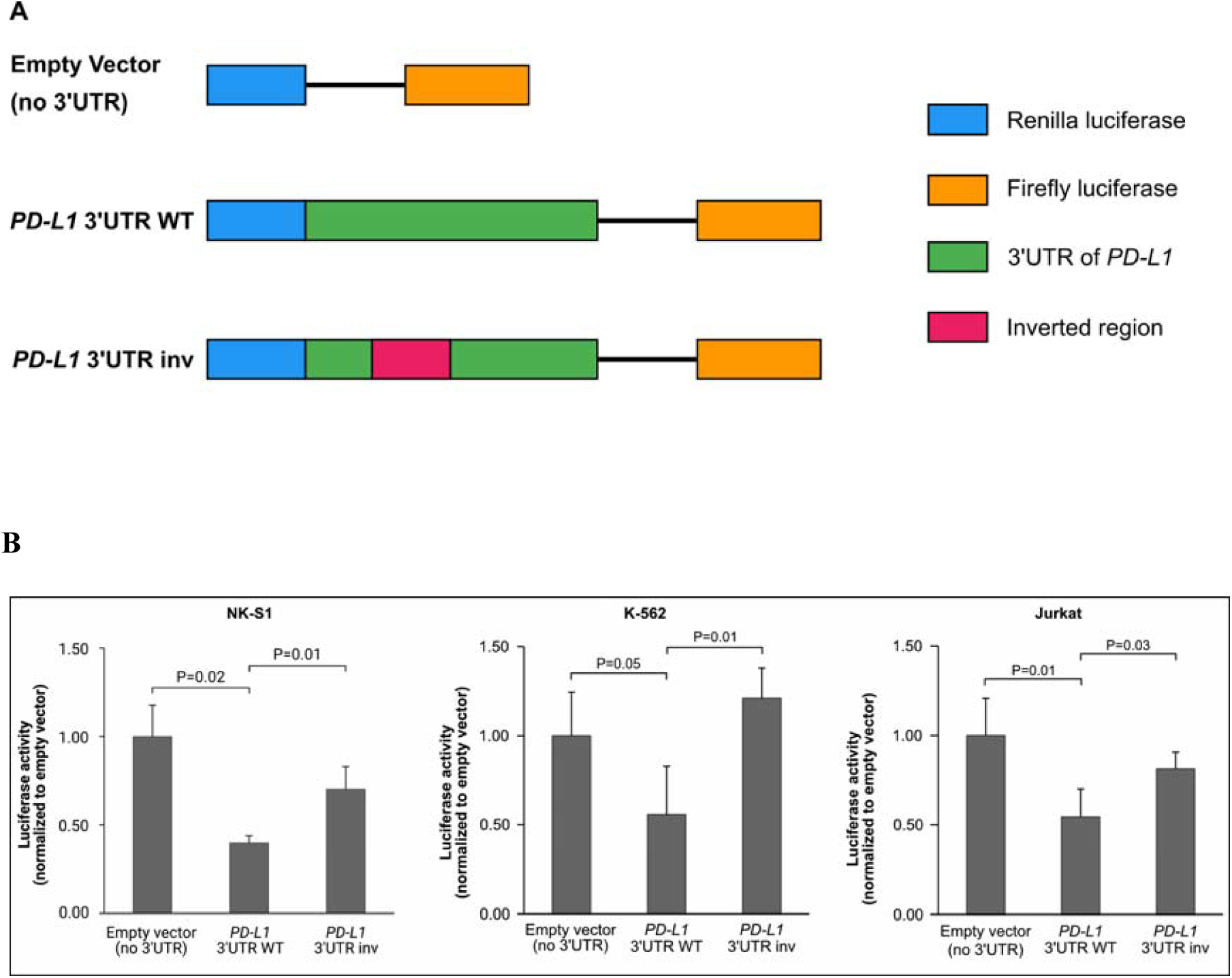
The effect of *PD-L1* 3’UTR on its expression. (A) The schematics of the used luciferase reporter constructs. Both full-length wild type (WT) and partially inverted (inv) *PD-L1* 3’UTR were cloned into the psiCHECK-2 vector. The partially inverted construct represents the rearrangement identified in NKTL1. Each element in the reporter is represented by a different color. (B) NK-S1, K-562 and Jurkat cells transiently transfected with WT or inv *PD-L1* 3’UTR reporter constructs respectively. Luminescence was measured after 48 h. Results are presented as a mean normalized to the empty vector control (mock). T bars represent standard deviations and *P* value was computed by two-sided t-test

Besides sequence analysis by our established genomic pipeline, visual inspection was also performed for known recurrent mutated genes of NKTL to avoid artefacts (*10-13, 16, 20, 21*). Among frequently mutated genes in NKTL, *TP53* stopgain (p.W14X) and *STAT3* missense (p.H410R) were also found in NKTL27 and NKTL28, respectively. Mutations in genes associated with antigen presentation and interferon gamma pathways, which are known to associate resistance to immune checkpoint blockade in melanoma (*22*), are not found in our cohort.

### Regulatory activity of *PD-L1* 3’UTR in NKTL

All four novel *PD-L1* SRs were predicted to lose whole or part of the *PD-L1* 3’UTR, except the micro-inversion that spanned across 206 bp and sits entirely within the 3’UTR of *PD-L1.* To determine the functional significance of this micro-inversion in regulating PD-L1 expression, the wild type and mutant (with 206 bp inversion) *PD-L1* 3’UTR were cloned into a luciferase reporter assay system and transfected into lymphoma and leukemia cell lines, namely, NK-S1, K-562 and Jurkat (Fig. 2A). Our results show that the wild type *PD-L1* 3’UTR could effectively suppress the luciferase activity of the reporter protein and the identified micro-inversion could relieve this suppression in NK-S1, K-562 and Jurkat cell lines (P = 0.01, P = 0.01 and P = 0.03, two-tailed t-test; Fig. 2B). Indeed, we have observed moderate to high levels (range: 20%-100%) of PD-L1 positivity in these four tumors harboring *PD-L1* 3’UTR SR (Table 2). These results offer a direct explanation to how these NKTL tumors would have evaded immune surveillance by upregulating PD-L1 expression.

### *PD-L1* structural rearrangements and *JAK3-*activating mutations are clonal in NKTL

Although the mechanisms of response to PD-1 blockade from *PD-L1* 3’UTR SRs and *JAK3*-activating mutations remain to be elucidated, we wanted to investigate if the clonality of these alterations could support the CR in patients who had *PD-L1* and *JAK3* alterations in their pretreated tumors, from the single-agent regime of pembrolizumab. From the somatic variants called, we were able to obtain solutions for the clonal architectures of 10 cases (SciClone (*23*) did not have a clonality solution for NKTL1). Five cases, four CR cases and one PD cases, had a clonal architecture (Table 3 and Supplementary Fig. 4). The somatic *PD-L1* and *JAK3* alterations identified in this study resided in the founding clone of their corresponding pretreated tumors. Given these results, the clonality analysis does support the extent of response in our patients who achieved CR from the single-agent regime of pembrolizumab therapy.

**Table 3.** Clonal residencies of the genomic correlates of response to pembrolizumab in the pretreated tumors of our study cohort CR, complete response; PD, progressive disease.

| **Sample ID** | **Response to Pembrolizumab** | **Number of Clones** | ***PD-L1* rearrangement clonal?** | ***JAK3*-activating mutation clonal?** |
| --- | --- | --- | --- | --- |
| NKTL1 | CR | no solution | Yes | - |
| NKTL26 | CR | 1 | Yes | - |
| NKTL28 | CR | 2 | Yes | - |
| NKTL31 | CR | 1 | Yes | - |
| NKTL29 | CR | 1 | - | Yes |
| NKTL30 | CR | 1 | - | Yes |
| NKTL43 | CR | 2 | - | - |
| NKTL25 | PD | 1 | - | - |
| NKTL27 | PD | 3 | - | - |
| NKTL44 | PD | 2 | - | - |
| NKTL45 | PD | 2 | - | - |

CR, complete response; PD, progressive disease.

## Discussion

Immunotherapy, in particular PD-1 blockade therapy, has shown promise in the treatment of several cancers, including NKTL. Recent study on adult T cell lymphoma (ATLL) has uncovered recurrent disruptions to the 3’UTR of *PD-L1* that induce tumor immune evasion via aberrant expression of PD-L1 (*24*). However, the study did not have concurrent clinical response data to further associate *PD-L1 3’UTR* SR and response to pembrolizumab. In our study, we showed that four out of seven (57%) who achieved CR to PD-1 blockade had clonal *PD-L1 3’UTR* SR and demonstrated that the SR could disrupt the suppression function of wild type *PD-L1* 3’UTR. *PD-L1* 3’UTR SR was also recently identified in a single case of ovarian cancer where the patient achieved CR with pembrolizumab *(25),* further supporting its role as a potential biomarker of response to PD-1 blockade therapy in NKTL.

All NKTLs are diagnostically EBER+ and the EBV protein, *LMP1* could constitutively upregulate PD-L1 (*26*). It is reasonable to speculate that NKTLs will respond to PD-1 inhibitors as they are innately PD-L1+. Indeed, all seven RR-NKTL patients in our previous clinical study had an initial response to pembrolizumab (*27*). However, at the time of this writing, three of the seven patients have progressed and died of disease. We did not find alterations of the *PD-L1* and *JAK3* genes in these PD cases. Conceivably, this initial “pseudo-remission” could be attributed by exogenous factors, such as EBV upregulating PD-L1 that was transiently blocked by initial dosages of pembrolizumab. Hence, high PD-L1 positivity in tumors will not necessarily equate to good response to PD-1 blockade. In addition, the PD-L1 IHC scores also varied greatly (6%, 2+ to 100%, 3+) within our cohort, and both NKTL25 and NKTL27 had PD despite having high PD-L1 staining grade for their pretreated tumors, questioning the effectiveness of PD-L1 positivity alone as a biomarker of response to PD-1 blockade in NKTL.

Interestingly, no rearrangements were identified within the *PD-L2* gene and *PD-L1* always served as the 5’ rearrangement partner. This is in contrast to other hematologic malignancies where the over-expression of PD-L1 and/or PD-L2 is achieved by diverse mechanisms such as genomic amplification, *JAK2* or *PD-L2* translocations (Supplementary Table 3) (*24, 28, 29*). This suggests that different tumors have evolved alternate mechanisms for immune evasion.

The presence of recurrent *JAK3-*activating mutations in our CR cases also coincides with a recent report showing the long-term benefit of PD-1 blockade in a single lung cancer patient with *JAK3-*activating mutations (*30*). Moreover, Zaretsky *et al* recently reported that loss of *JAK1*, *JAK2* or *B2M*, leading to a lack of response to interferon gamma or antigen presentation confers resistance to pembrolizumab in metastatic melanoma (*22*). These data led us to speculate that *JAK3-*activating mutations could play an important role in patient’s response to immune checkpoint inhibitors and the mechanisms require further investigation.

The initial tumor from one of our CR patients (ID: NKTL43), both *PD-L1*^WT^ and *JAK3*^WT^, harbored a 3 bp insertion in the *ARID1B* gene (Supplementary Fig. 5). A recent study has reported PBRM1-deficient and ARID2-deficient tumors correlated with better response to anti-PD-1/PD-L1 therapy renal cell carcinoma (*31*). There seems to be a relationship between truncating alterations in the subunits of the SWI/SNF complex and response to PD-1/PD-L1 therapy. However, the mechanisms and which members of these large complexes are involved in conferring response to immune checkpoint blockade therapy await further investigation.

To the best of our knowledge, this is the first study to report on the correlates of genomic features with response to PD-1 blockade therapy in NKTL using WGS data. The main limitation of this study is the small sample size and this is mainly due to the rarity of NKTL and also that PD-1 blockade is still not approved for the treatment of RR NKTL. In summary, PD-1 blockade in NKTL is promising and genomic correlates of response from our study will have to be further validated in larger cohorts of NKTL patients treated with immune checkpoint therapy.

## Materials and Methods

### Patients and study setup

Our study cohort consists of 11 patients with RR NKTL who had failed L-asparaginase-based chemotherapy regimens from Singapore, China and Hong Kong. NKTL1, NKTL25, NKTL26, NKTL43, NKTL44 and NKTL45 were never sequenced before and were included from our previous study (*27*). Patients were diagnosed with NKTL according to the 2008 World Health Organization classification with cytotoxic, CD3ε+ and EBER+ phenotypes (*3*). Response assessment was performed using a combination of PET/CT or CT/MRI, EBV PCR, and in one patient, histological assessment of a resected lesion that was fluorodeoxyglucose avid on PET/CT scan. The duration of response (DoR) was calculated from the date of starting pembrolizumab to the date of progression or death. The median DoR was estimated using the Kaplan-Meier method. Institutional Review Boards from SingHealth (2004/407/F), National University of Singapore (NUS-IRB-10-250) and Sun Yat-sen University Cancer Center (YB2015-015-01) approved the study. All subjects in this study provided written informed consent. The study also adhered to the Declaration of Helsinki.

### Genomic DNA extraction

Genomic DNA from snap frozen and formalin-fixed paraffin-embedded (FFPE) tumor tissues, and whole blood was extracted as previously described (*32*). Buccal swab genomic DNA was purified using E.Z.N.A. Tissue DNA Kit (Omega Bio-tek) according to manufacturer’s instructions. The quality and quantity were assessed as described elsewhere (*32*). All sequenced cases in our study, except NKTL25 being a snap frozen sample, were FFPE samples.

### Whole genome sequencing

Whole-genome sequencing (WGS) was performed for all 11 pairs of tumor-normal samples described in this study. All sequencing libraries were prepared using TruSeq Nano DNA Library Prep Kit (Illumina). Due to high fragmentation of genomic DNA from FFPE material, a size selection step was conducted prior to library preparation for the FFPE tumor samples. Amplifiable DNA fragments of ∼200 bp from our FFPE samples are used for sequencing library construction to avoid false-negatives confidently in our discovery for SR within the *PD-L1* gene. The gDNA from the FFPE tumors were sequenced on the HiSeq4000 System (Illumina) with a 2×150 bp format and the gDNA for the matching normals were sequenced on the HiSeq X System (Illumina) with a 2×150 bp format.

### Detection and filtering of structural rearrangements

Sequencing reads were aligned using BWA-MEM (*33*) to the hs37d5 human reference genome. Strelka2 (*34*) and MuSE (*35*) were used to detect somatic short variants. Variants were subsequently annotated by wAnnovar (*36*) on 7^th^ August 2018. The genic regions of *PD-L1* and *PD-L2* were manually inspected for somatic structural rearrangements with IGV. Read pairs were marked as discordant if they did not align to the reference genome with the expected orientation and/or within the expected insert size. Reads were flagged as clipped when either end of the read did not match the reference genome. Strelka2 was run with default WGS profile. MUSE was run with default parameters.

### Analysis of tumor clonality

SciClone (*23*) was used to analyze the clonality architecture of the tumors. CANVAS (*37*) was used to analyze copy number and loss of heterozygousity information for each tumor, which were used as input for the clonality analysis by SciClone.

### PCR and Sanger sequencing

Details about PCR conditions and sequencing are described elsewhere (*38*). Primers were designed using Primer3 software (*39*) and the sequences are listed in Supplementary Table 4. Sanger sequences were aligned to human reference genome and confirmed with BLAT (*40*).

### Histological studies and scoring

PD-L1 IHC analysis was performed with anti-PD-L1 rabbit monoclonal antibody (SP263, Ventana) by Dr Jabed Iqbal. PD-L1 positivity was evaluated as a percentage of positively stained tumor cells at the cell membrane.

### Cell lines and constructs

K-562 and Jurkat cell line was purchased from ATCC and NK-S1 (*41*) was generated in-house. LGC Standards authenticated the K-562 and Jurkat cell lines on 6^th^ October 2017. Jurkat cells were maintained in RPMI 1640 (Gibco) supplemented with 10% FBS (HyClone), and K-562 and NK-S1 were grown in DMEM (Gibco) supplemented with 10% FBS (HyClone), 10% horse serum (Gibco) and 2 mM L-glutamine (Gibco). The cells were grown at 37°C in the presence of 5% CO_2_ and routinely checked for mycoplasma contamination using MycoAlert Mycoplasma Detection Kit (Lonza).

Wild type *PD-L1* 3’UTR (ENST00000381573.8) from SNK6 cell line (generously provided by Dr. Norio Shimuzu) was cloned into the XhoI and NotI sites of the psiCHECK-2 vector (Promega). For the partially inverted 3’UTR recapitulating the rearrangement identified in sample NKTL1, three individual pieces with overhangs were amplified from a wild type sample (SNK6) and ligated together by PCR. Cloning was performed using Q5 High-Fidelity 2X Master Mix (New England BioLabs). All cloning primers used to clone the full-length wild type and mutant *PD-L1* 3’UTR are described in Supplementary Table 5.

### Transfection and luciferase assay

For K-562 and Jurkat, 5×10^4^ cells and 6×10^4^ cells were seeded on a 48-well plate in triplicates, respectively, and transfected with 250 ng plasmid DNA using the Lipofectamine 3000 Reagent (Invitrogen). For NK-S1 cells, 2×10^5^ cells were electroporated in triplicates on a 24-well plate with 1 µg plasmid DNA using the Neon Transfection System (Invitrogen). The pulse parameters used were the following: voltage 1300, width 10 and no. 3.

The cells were lysed with Passive Lysis Buffer (Promega) after 48 hours. Luminescence was measured using the Dual-Luciferase Reporter Assay System (Promega) and the GloMax-Multi+ Detection System (Promega). Renilla luciferase activities were divided by Firefly luciferase activities and the results were normalized to the empty vector control (mock). Statistical significance was calculated by two-sided t-test. Statistical significance was considered as P<0.05. All experiments were repeated at least twice.

## Data availability

The WGS data of 11 ENKL-normal/blood pairs have been deposited in European Genome-phenome Archive (EGA) under the study accession code: EGAS00001002420.

## List of Supplementary Materials

Supplementary Fig. 1. Recurrent somatic mutated genes in the 11 pembrolizumab-treated patients’ initial tumors.

Supplementary Fig. 2. Validation of *PD-L1* rearrangements and *JAK3*-activating identified in Natural-Killer/T-Cell Lymphoma patients with complete response to pembrolizumab.

Supplementary Fig. 3. Schematics of the tandem duplication disrupting the 3’UTR of *PD-L1* in NKTL26 inferred from whole genome sequencing data.

Supplementary Fig. 4. Clonality cluster plots from SciClone.

Supplementary Fig. 5. Validation of the 3bp insertion in *ARID1B* Identified in a Natural-Killer/T-Cell Lymphoma patient with complete response to pembrolizumab.

Supplementary Table 1. Statistics of the whole-genome sequencing data in our study.

Supplementary Table 2. Predicted non-silent protein-coding mutations.

Supplementary Table 3. *PD-L1* and *PD-L2* alterations described in hematological malignancies.

Supplementary Table 4. Primer-pairs used for the validation of *PD-L1* structural rearrangement, *JAK3*-activating and *ARID1B* mutations.

Supplementary Table 5. Cloning primers used for the cloning of the full-length 3’UTR of *PD-L1* with and without the micro-inversion of 206 bp long.

